# A fungal RNA-dependent RNA polymerase is a novel player in plant infection and cross-kingdom RNA interference

**DOI:** 10.1101/2023.06.02.543307

**Authors:** An-Po Cheng, Bernhard Lederer, Lorenz Oberkofler, Lihong Huang, Fabian Platten, Florian Dunker, Constance Tisserant, Arne Weiberg

**Affiliations:** Institute of Genetics, Faculty of Biology, Ludwig Maximilian University of Munich, 82152 Martinsried, Germany; GMU-GIBH Joint School of Life Sciences, The Guangdong-Hong Kong-Macau Joint Laboratory for Cell Fate Regulation and Diseases, Guangzhou Medical University

**Author notes:** Correspondence: Arne Weiberg.

## Abstract

Small RNAs act as fungal pathogen effectors that silence host target genes to promote infection, a virulence mechanism termed cross-kingdom RNA interference (RNAi). The essential pathogen factors of cross-kingdom small RNA production are largely unknown. We here characterized the RNA-dependent RNA polymerase (RDR)1 in the fungal plant pathogen *Botrytis cinerea* that is required for pathogenicity and cross-kingdom RNAi. *B. cinerea bcrdr1* knockout (ko) mutants exhibited reduced pathogenicity and loss of cross-kingdom small RNAs. We developed a novel “switch-on” GFP reporter to study cross-kingdom RNAi in real-time within the living plant tissue which highlighted that *bcrdr1* ko mutants were compromised in cross-kingdom RNAi. Moreover, blocking seven pathogen cross-kingdom small RNAs by expressing a short-tandem target mimic RNA in transgenic *Arabidopsis thaliana* led to reduced infection levels of the fungal pathogen *B. cinerea* and the oomycete pathogen *Hyaloperonospora arabidopsidis*. These results demonstrate that cross-kingdom RNAi is significant to promote host infection and making pathogen small RNAs an effective target for crop protection.

## Introduction

RNA-dependent RNA polymerases (RdRPs, RDRs) synthesize a complementary RNA strand from a primary RNA template without requiring any free DNA or RNA primer. Two classes of RdRPs/RDRs have been described. The viral-type RdRPs have been well-characterized in viral replication and have been also found as parts of retrotransposon genomes. Viral-type RdRPs are promising therapeutic targets to cure COVID-19 and other viral infections (Machitani *et al*., 2020). The cellular-type RDRs are conserved in several eukaryote kingdoms, including plants, nematodes, and fungi, and are functional in RNA interference (RNAi) pathways (Chang *et al*., 2012, Mello & Conte, 2004). RDRs generate double-stranded RNA precursors for the DICER or Dicer-like (DCL)-dependent biogenesis of primary and secondary small-interfering (si)RNAs that load into Argonaute (AGO) proteins and form an RNA-induced silencing complex (RISC) inducing transcriptional or post-transcriptional gene silencing.

In the model plant *Arabidopsis thaliana*, the RDR2 synthesizes from DNA polymerase IV-dependent RNA transcripts double-stranded RNAs (dsRNAs) that initiate small RNA-directed DNA methylation (RdDM). RDR2 and the RdDM pathway are mainly responsible for the transcriptional control of transposons and are involved in transgene-induced gene silencing (Matzke & Mosher, 2014). The *A. thaliana* RDR6 is required to produce secondary siRNAs, an amplification loop that fosters gene silencing of RNA virus and transgenes, as well as controlling endogenous gene expression. Plant RDR2 and RDR6 are both involved in the molecular stress response and defense against various types of plant pathogens, including virus, bacteria, fungi, and oomycetes (Boccara *et al*., 2014, Garcia-Ruiz *et al*., 2010). RDRs are also required for RNAi and the biogenesis of various types of small RNAs in fungi (Torres-Martinez & Ruiz-Vazquez, 2017). Two distinct RDR-dependent RNAi pathways have been elucidated in the fungus *Neurospora crassa*. The RDR QDE-1 is part of the quelling pathway, an RNAi mechanism that controls transgene-induced gene silencing. A second *N. crassa* RDR, SAD-1, produces dsRNAs to induce meiotic silencing of unpaired DNA (MSUD), controlling aberrant RNA accumulation during late meiosis (Chang *et al*., 2012). Two distinct RDRs of the basal fungus *Mucor circinelloides* are required for initiation and amplification of transgene-induced gene silencing (Calo *et al*., 2012). Like plants, fungal RDRs in the species *Cryphonectria parasitica* are functional in antiviral defense and transposon silencing (Zhang *et al*., 2014); however, a RdDM pathway as known in plants has not been found in fungi. Moreover, RDRs have been proposed to regulate genes affecting fungal growth and development, but little is known about the functional role of fungal RDRs in pathogenicity. A knockout of *MoRdRP1* in the rice blast fungus *Magnaporthe oryzae* led to reduced growth, conidia formation, and disease symptoms on rice leaves (Raman *et al*., 2017). Similarly, *fgrdrp2, fgrdrp3* and *fgrdrp4* knockout strains of the head blight inducing fungal pathogen *Fusarium graminearum* indicated reduced conidia, mycotoxin production, and head blight on inoculated wheat spikes (Gaffar *et al*., 2019). While such reports have indicated that distinct fungal RDRs are involved in pathogenicity, their mode of action remains unclear.

In this study, we investigated the role of the *BcRDR1* gene in the fungus *Botrytis cinerea*, a multi-host plant pathogen infecting more than 1,000 different plant species and causing the devastating grey mold disease. As part of its pathogenicity, *B. cinerea* releases extracellular small RNAs (Bc-sRNAs) that enter plant cells during infection and recruit the plant’s own AGO1/RISC to induce host gene silencing for promoting infection (Wang *et al*., 2017, Weiberg *et al*., 2013). This virulence phenomenon has been termed cross-kingdom RNAi and is a general infection strategy of diverse plant and animal pathogenic as well as symbiotic microbes, including fungi, oomycetes, and bacteria (Cai *et al*. 2019, Weiberg *et al*., 2015, Weiberg *et al*., 2014). Most Bc-sRNAs inducing cross-kingdom RNAi are encoded by GYPSY-class of long terminal repeat retrotransposons and are produced by *B. cinerea* BcDCL1 and BcDCL2 (Porquier *et al*., 2015, Weiberg *et al*., 2014, Weiberg *et al*., 2013). We herein discovered that the *B. cinerea* BcRDR1 is a pathogenicity factor and is required for *B. cinerea*-induced cross-kingdom RNAi.

## Results

### *B. cinerea* BcRDR1 is a pathogenicity factor

In order to identify unknown genes involved in cross-kingdom RNAi, were analyzed the family of BcRDRs in *B. cinerea*. Through a genome survey of the *B. cinerea* strain B05.10, we identified three genes encoding conserved RDRs, thereafter named BcRDR1 (Bcin01g01810), BcRDR2 (Bcin08g02220), and BcRDR3 (Bcin07g01750). Phylogenetic analysis of full-length amino acid sequences revealed that BcRDR1 is the orthologue of the *N. crassa* SAD-1 involved in MSUD (Shiu *et al*. 2001), and BcRDR2 is the orthologue of *N. crassa* quelling RDR, QDE-1 (Cogoni & Macino 1999). BcRDR3 is another conserved RDR with unknown function (Figure S1A). All three BcRDRs comprised a conserved DxDGD motif indicating that they function as RNA polymerases (Figure S1B). We detected transcripts of *BcRDR1, BcRDR2* and *BcRDR3* when growing *B. cinerea* in liquid culture or during infection of *Solanum lycopersicum* (tomato) leaves; however, there was no significant up-regulation of any of the *BcRDR*s measured during infection (Figure S2A).

In order to investigate potential functions of BcRDRs in *B. cinerea* pathogenicity, we generated targeted gene knockout (ko) mutant strains by homologous recombination and obtained two independent *bcrdr1* ko isolates (Figure S2B). Infection assays using detached *S. lycopersicum* or *A. thaliana* leaves revealed that *bcrdr1* ko mutants induced smaller lesion sizes compared to the *B. cinerea* wild type (WT) strain, which was accompanied with lower fungal biomass, as estimated by quantification of *B. cinerea Tubulin* mRNAs levels (Figure 1A-C and Figure S3). Both *bcrdr1* ko mutants showed a WT-like growth and development phenotype under axenic culture condition (Figure 1D and E). Based on these results, we assume that BcRDR1 is a pathogenicity factor in *B. cinerea*.

**Figure 1:**
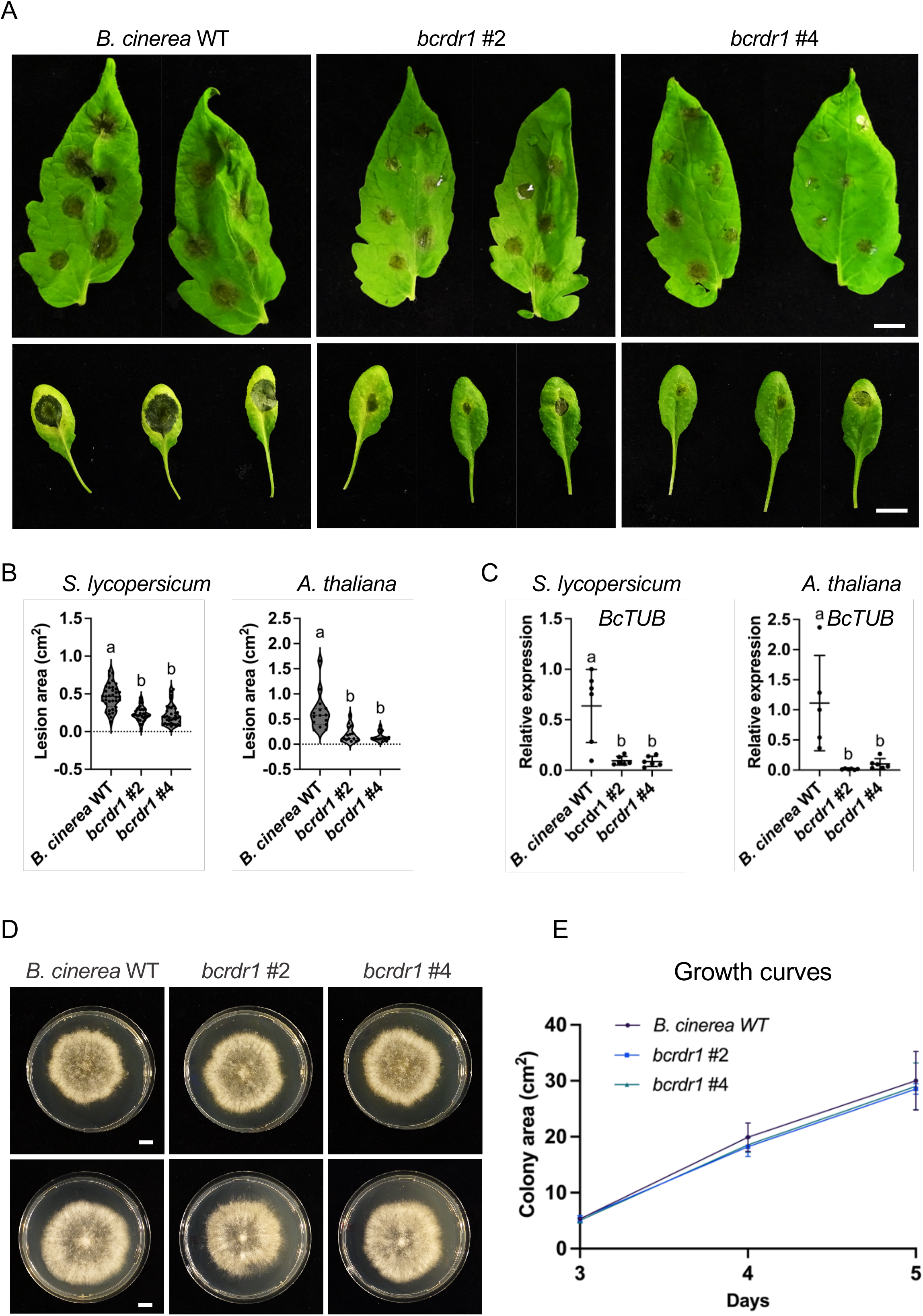
*B. cinerea bcrdr1* ko mutants are compromised in pathogenicity. A) Infection series on *S. lycopersicum* or *A. thaliana* detached leaves using *B. cinerea* WT and two *bcrdr1* ko mutants #2 and #4. B) Lesion size induced by *B. cinerea* infection was measured at 48 hpi. A minimum of 34 lesions on *S. lycopersicum* and 15 lesions on *A. thaliana* were measured and statistical analysis was performed using ANOVA followed by a Tukey post-hoc test with *p*-value threshold *p* < 0.05. The scale bars represent 1 cm. C) *B. cinerea* biomass was estimated by measuring mRNA levels of the reference gene *Bc-Tubulin (BcTUB)* and related to the plant reference *SlActin* or *AtActin* in four biological replicates. Statistical analysis was performed using ANOVA followed by a Tukey post-hoc test with *p*-value threshold *p* < 0.05. The scale bars represent 1 cm. D) Colony grow of B. cinerea WT and *bcrdr1* mutants #2 and #4 on malt extract agar after 5 days. The scale bars represent 1 cm. E) Growth curves of *B. cinerea* WT and *bcrdr1* ko mutants was measured in 5 replicates over a time course of 5 days. Error bars indicate the standard deviation of the 5 replicates.

### Biogenesis of retrotransposon-derived Bc-sRNAs requires BcRDR1

Some plant, nematode, and fungal RDRs are required for siRNA biogenesis, and stopping Bc-sRNA production in a *B. cinerea bcdcl1dcl2* ko mutant leads to reduced pathogenicity due to impaired cross-kingdom RNAi (Weiberg *et al*., 2013). To investigate the role of BcRDR1 in *B. cinerea* Bc-sRNA biogenesis, we performed comparative small RNA-seq analysis. *B. cinerea* WT and two independent *bcrdr1* ko mutants were cultured for two days in liquid medium, and mycelium was collected to extract total RNA. Upon small RNA isolation via PAGE, RNA libraries were cloned for Illumina sequencing. In total, 1.3 – 4.9 million reads in depth were mapped to the *B. cinerea* reference genome either unique or multiple times. The fraction of multiple mapping reads was reduced in both *bcrdr1* ko mutants (Figure 2A). Plotting read numbers by size revealed a reduction of 21-22 nucleotides (nt) long Bc-sRNAs in both *bcrdr1* ko mutants, mostly representing 5’ prime Uracil nucleobase, retrotransposon (RT)-derived Bc-sRNAs (Figure 2B-D). This result indicated that production of Bc-sRNAs that mostly mapped to RTs was dependent on BcRDR1. The long-terminal repeat RT *BcGypsy3* is a major source of Bc-sRNAs that are induced during plant infection, and is a pathogenicity factor of *B. cinerea* (Porquier *et al*., 2021). Accordingly, reduced accumulation of RT-derived Bc-sRNAs in *bcrdr*1 ko mutants resulted in increased mRNA levels of *BcGypsy3* (Figure S4), suggesting a role of BcRDR1 in post-transcriptional silencing of RTs.

**Figure 2:**
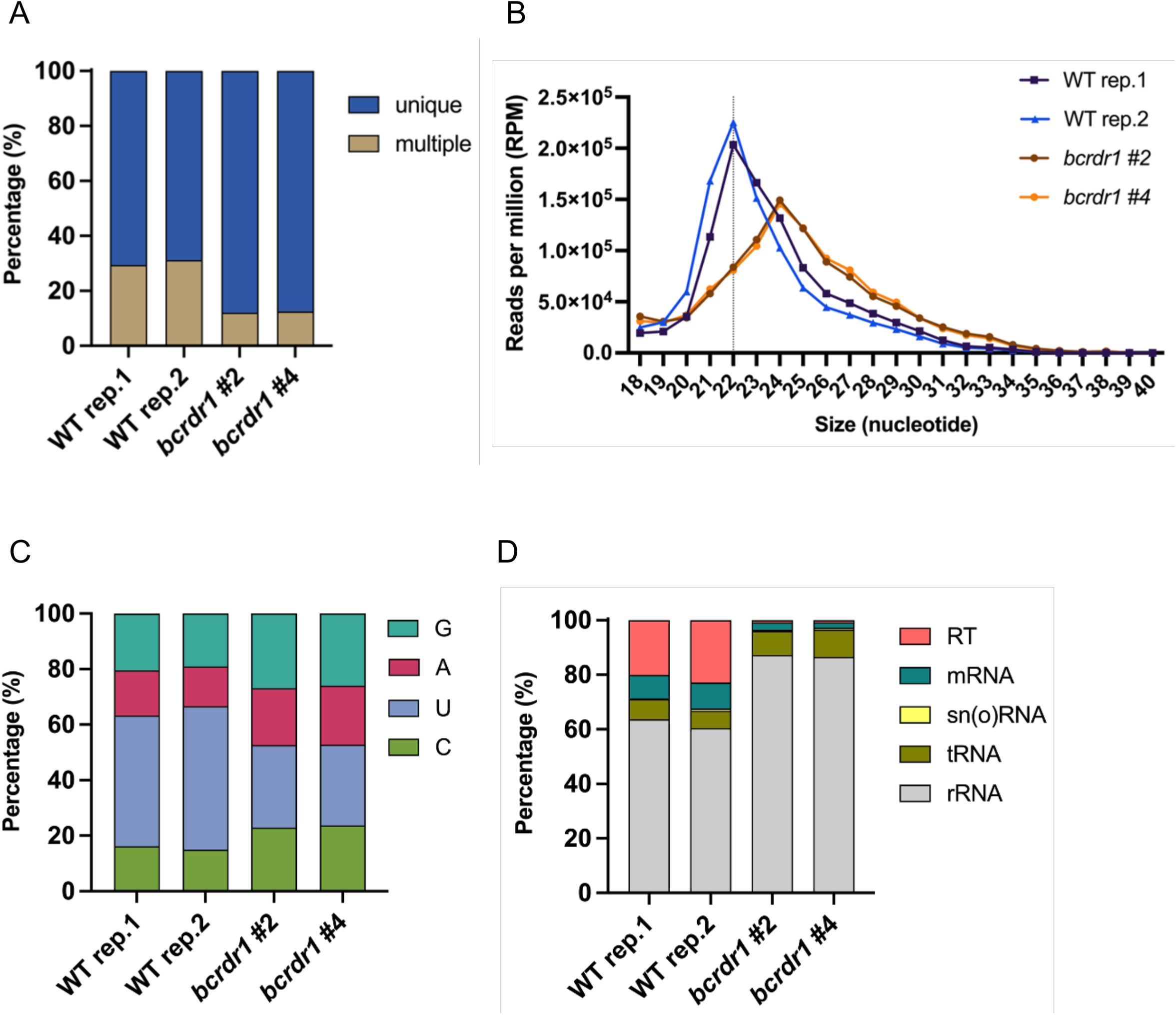
Small RNA sequencing analysis of *B. cinerea* WT and *bcrdr1* ko mutants. A) Fractions of Bc-sRNAs mapping unique or multiple times to a *B. cinerea* reference genome. B) Bc-sRNA size profiles (18-40 nt) of mapped Bc-sRNA reads in reads per million (RPM).C) Distribution in percentage of the four RNA nucleotides C, U, A, G at the 5’prime position of Bc-sRNAs. D) Distribution in percentage of Bc-sRNAs mapping to distinct *B. cinerea* RNA gene loci: ribosomal RNA (rRNA), transfer RNA (tRNA), small nuclear and nucleolar RNA (sn(o)RNA), messenger RNA (mRNA), and retrotransposon (RT).

### *B. cinerea bcrdr1* ko mutants are compromised in cross-kingdom RNAi

RT-derived Bc-sRNAs were previously found to induce cross-kingdom RNAi (Porquier *et al*., 2021, Weiberg *et al*., 2013). In particular, the four Bc-sRNAs, Bc-sRNA3.1, Bc-sRNA3.2, Bc-sRNA5, and Bc-sRNA20 induced silencing of the *S. lycopersicum* host genes *Valuolar sorting protein* (*SlVPS*) (Solyc09g014790), *Mitogen-activated protein kinase kinase kinase* (*SlMPKKK)4* (Solyc08g081210), and *C2H2 zinc-finger transcription factor SlBhlh63* (Solyc03g120530), and the *A. thaliana* host genes *AtMPK1* (AT1G10210), *AtMPK2* (AT1G59580), a *Cell Wall kinase* (*WAK*) (AT5G50290), and the *Peroxredoxin PRXIIF* (AT3G06050) (Figure 3A). We confirmed that *bcrdr1* ko mutants lost accumulation of these Bc-sRNAs by stem-loop reverse transcriptase PCR (Figure 3B). When infecting *S. lycopersicum* or *A. thaliana* with *B. cinerea bcrdr1* ko mutants, down-regulation of Bc-sRNA target genes, as observed in WT-infected plants infected with the wild-type strain compared to non-inoculated plants, was abolished or less strong (Figure 3C-D, Figure S5A-B). Successful infection of plants and induced host defense response due to *B. cinerea* infection was validated by measured up-regulation of *S. lycopersicum Sl Proteinase inhibitor* (*PI*)*-I* and *SlPI-II* or *A. thaliana AtPlant Defensin* (*PDF*)*1*.*2* immunity genes. These results indicated that *bcrdr1* mutants might be compromised in inducing cross-kingdom RNAi, which would explain the reduced pathogenicity observed in *S. lycopersicum* and *A. thaliana*.

**Figure 3:**
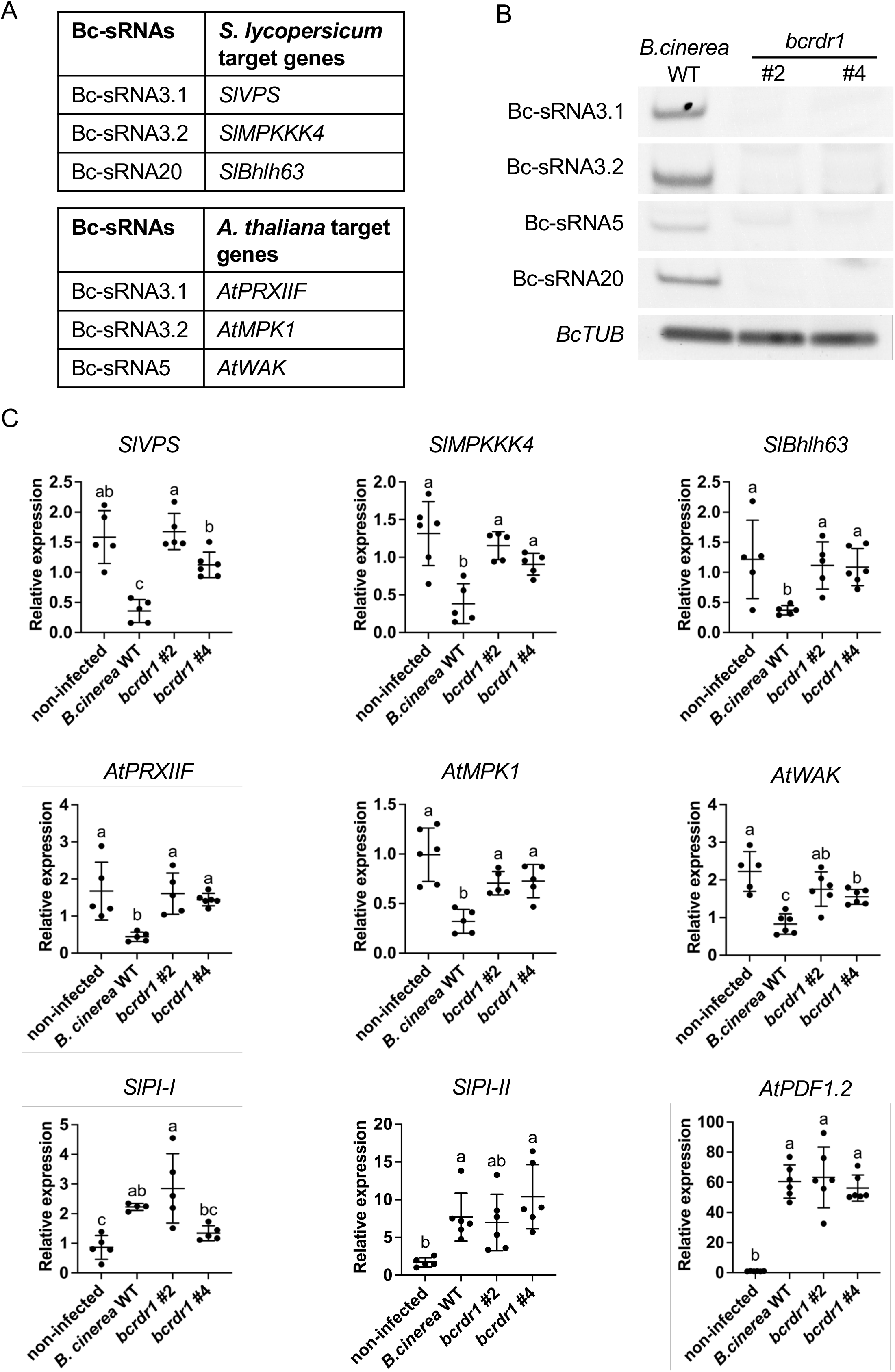
*B. cinerea bcrdr1* ko mutants are compromised in plant target gene suppression. A) Known *S. lycopersicum* and *A. thaliana* target genes silenced by Bc-sRNAs through cross-kingdom RNAi. B) Stem-loop reverse transcriptase PCR of Bc-sRNAs in *B. cinerea* WT and *bcrdr1* ko mutants. *BcTUB* was used as internal control. C) Quantitative reverse transcriptase PCR measuring mRNA levels of Bc-sRNA target genes in *S. lycopersicum* and *A. thaliana* during infection with *B. cinerea* WT or *bcrdr1* ko mutants. Lines in scatter plots represent the mean and the standard deviation. Each gene was measured in at least five biological replicates. *Sl-Actin* or *At-Actin* were used as reference genes. *SlPI-I, SlPI-II* and *AtPDF1*.*2* were measured as plant immunity marker genes induced during *B. cinerea* infection. Statistical analysis was performed using ANOVA followed by a Tukey post-hoc test with *p*-value threshold *p* < 0.05.

To further inspect whether *bcrdr1* mutant strains were indeed compromised in inducing cross-kingdom RNAi, we designed a novel GFP “switch-on” cross-kingdom RNAi reporter that exclusively responded to the translocation of Bc-sRNA3.1 and Bc-sRNA3.2 from *B. cinerea* into the plant host (Figure 4A). In this reporter construct, the CRISPR-type RNA endonuclease Csy4 (Haurwitz *et al*. 2010) is co-expressed with a GFP version that is fused to the Csy4 recognition motif at its N-terminus. We fused the native target sites of Bc-siRNA3.1 and Bc-siR3.2 to the 5’ prime or 3’ ends of the *Csy4* transgene, respectively, turning *Csy4* into a target gene of these Bc-sRNAs. Csy4 constantly suppresses expression of the *GFP*, unless GFP expression is activated when Bc-sRNAs silence *Csy4*. This GFP switch-on reporter was stably expressed in transgenic *A. thaliana* plants. With these reporter plants, we were able to visualize *B. cinerea*-induced cross-kingdom RNAi in infected leaf tissue by fluorescent microscopy. Inoculating leaves of reporter plants with *B. cinerea* WT conidia suspension led to enhanced GFP expression compared to leaves inoculated with the *bcrdr1* #2 ko mutant (Figure 4B). The non-invasive GFP switch-on reporter allowed us to visualize cross-kingdom RNAi by life-time imaging. We recorded a time lapse video of GFP activity in reporter plants either inoculated with *B. cinerea* WT or *bcrdr1* ko #2 for 45 hours (Video 1). A clear increase in the GFP signal was apparent in *B. cinerea* WT infected leaves after 24 hpi (Figure 4C). We confirmed in independent infection series the activation of the GFP signal in plants infected with B. cinerea WT, but not in leaves inoculated with the two *bcrdr1* ko #2 and #4 mutants as well as when infected with the previously characterized *bcdcl1/bcdcl2* double knock-out (*bcdcl1/2*) mutant strain, which is unable to produce reporter-activating Bc-sRNAs (Weiberg *et al*., 2013) (Figure S6). We verified the results obtained by fluorescence microscopy when measuring GFP mRNA and protein levels by quantitative reverse transcriptase PCR and Western blot analysis using an anti-GFP antibody (Figure 4D and 4E). Referring to earlier in this study, we had observed that *bcrdr1* mutants developed less biomass during infection due to reduced virulence (Figure 1A). To rule out that reduced *B. cinerea bcrdr1* biomass could be responsible for lower GFP activity when inoculating the cross-kingdom RNAi reporter plants, we used 10x higher conidia suspension compared to *B. cinerea* WT inoculation and found that with increased conidia concentration, GFP activity was still significantly lower with *bcrdr1* ko (Figure S7). GFP activity was strongest at the infection front of *B. cinerea* WT inoculation sites (Figure S7B). This might indicate that cross-kingdom RNAi was the strongest in newly infected leaf cells.

**Figure 4:**
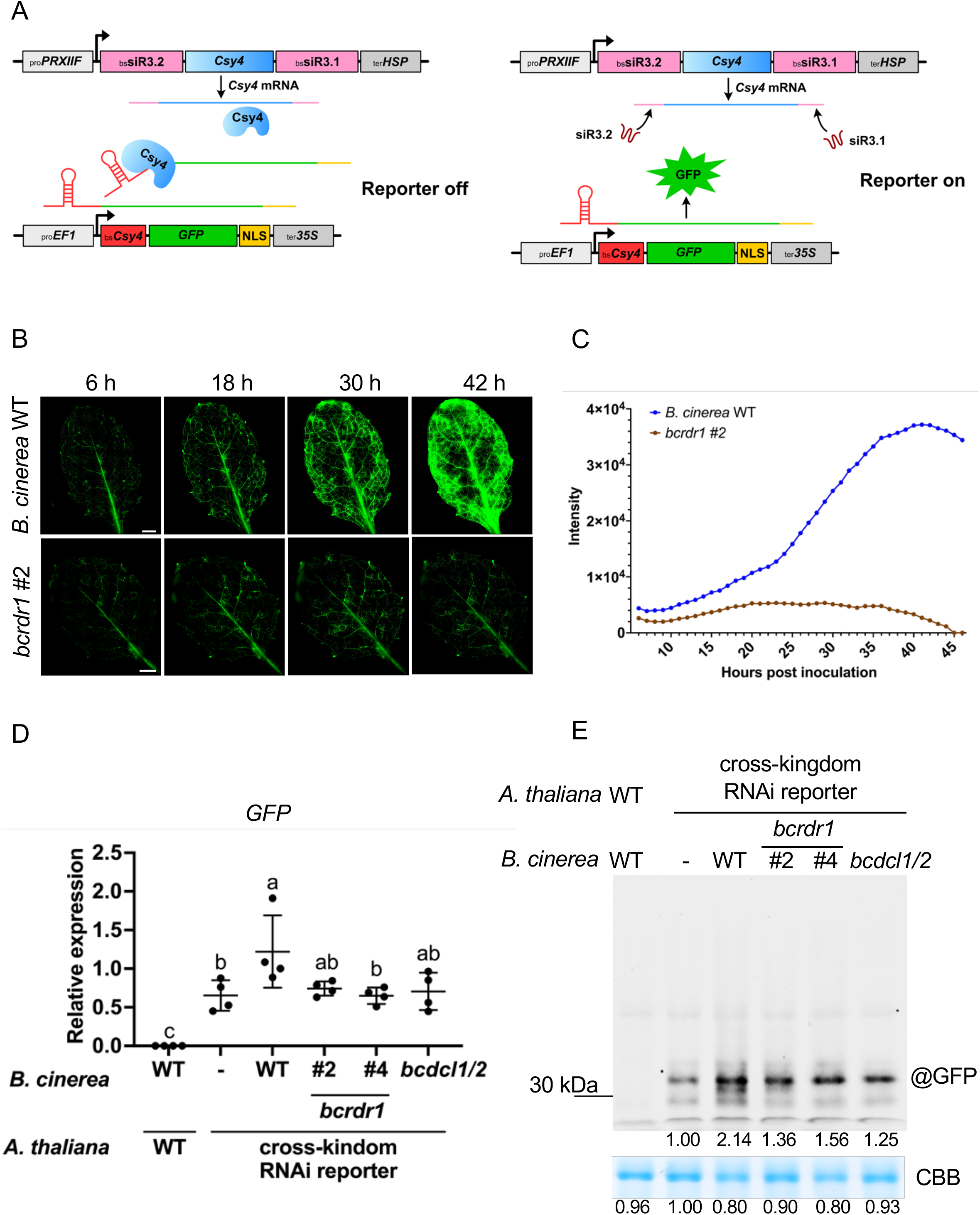
*B. cinerea bcrdr1* ko mutants are compromised in cross-kingdom RNAi. A) Schematic overview of a GFP-based switch-on cross-kingdom RNAi reporter suitable for *in planta* expression. B) GFP images from different time points. The scale bars represent 1 mm. C) GFP quantification in WT versus *bcrdr1* ko #2 from 6 – 46 hpi. D) Quantitative reverse transcriptase PCR analysis of relative *GFP* mRNA expression in cross-kingdom RNAi reporter plants during *B. cinerea* infection. Statistical analysis was performed using ANOVA followed by a Tukey post-hoc test with *p*-value threshold *p* < 0.05. The “-” symbol represents non-infected samples inoculated with water. E) Western blot analysis of GFP expression in cross-kingdom RNAi reporter plants during *B. cinerea* infection. The ribulose 1,5-bisphosphate carboxylase/oxygenase (RuBisCo) signal detected by Coomassie Brilliant Blue (CBB) staining was used as a loading control. Numbers indicate GFP and RuBisCo signal intensities estimated by the FIJI software. Samples in E) and F) were taken at 24 hpi.

### *B. cinerea* small RNAs translocated into plants promote infection

Several independent studies revealed that knockout or knockdown of DCLs led to loss of Bc-sRNA biogenesis and reduced pathogenicity in *B. cinerea* and other fungal pathogens (Gaffar *et al*., 2019, Qiao *et al*., 2021, Qiao *et al*., 2023, Wang & Jin, 2017, Wang *et al*., 2016, Weiberg *et al*., 2013, Werner *et al*., 2020, Werner *et al*., 2021, Zanini *et al*., 2021). Here, we demonstrate that knocking out the *BcRDR1* leads to loss of Bc-sRNA production, compromises cross-kingdom RNAi, and reduces pathogenicity. However, *BcRDR1* deletion might have affected other endogenous small RNA regulatory processes in the fungus that are relevant for infection (Göhre & Weiberg, 2022). Therefore, we aimed to further validate that cross-kingdom RNAi was part of *B. cinerea* pathogenicity. Transgenes producing small RNA sponges, for example RNA short tandem target mimic (STTM), can block microRNA- and siRNA-induced silencing of plant endogenous and exogenous target genes (Tang *et al*., 2012, Dunker *et al*., 2020). We cloned a Bc-siRNA3.2/Bc-sRNA5 double STTM (Figure 5A) and transformed it into *A. thaliana* for stable expression. We could isolate three independent *A. thaliana* STTM T2 lines and infected those lines with *B. cinerea* WT. Bc-sRNAs STTM plants exhibited reduced lesion sizes induced by *B. cinerea*, compared to *A. thaliana* WT or a transgenic *A. thaliana* line expressing a non-sense RNA-STTM (Figure 5B and Figure S8A). In consistence, Bc-sRNA3.2 and Bc-sRNA5 target genes in *A. thaliana, AtMPK1, AtMPK2*, and *AtWAK* were suppressed in *A. thaliana* WT or *A. thaliana* expressing a non-sense RNA STTM upon *B. cinerea* infection, but suppression of those host target genes was abolished in the Bc-sRNA STTM plant lines (Figure 5C and Figure S8B). Successful infection of all plant lines resulted in an induced host defense response due to *B. cinerea* infection which was validated by measured up-regulation of the *A. thaliana* immunity-related genes *AtBAK1* and *AtPDF1*.*2*. These results confirmed that cross-kingdom RNAi is an important part of *B. cinerea* pathogenicity.

**Figure 5:**
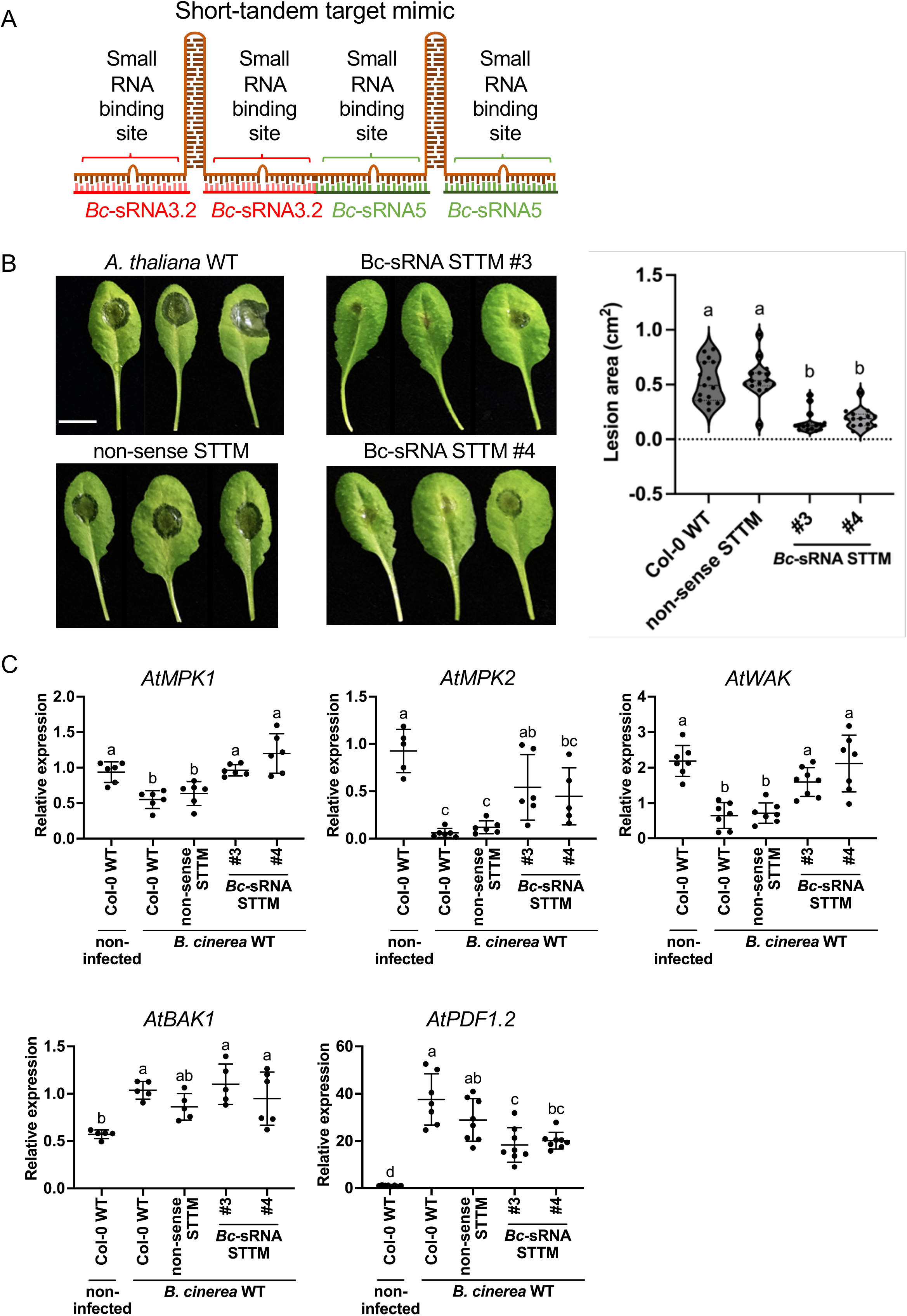
A plant expressing Bc-sRNA STTM blocks cross-kingdom RNAi and reduces *B. cinerea* pathogenicity. A) STTM construct to block Bc-sRNA3.2 and Bc-sRNA5 *in planta*. B) *A. thaliana* detached leaf inoculation assay using *B. cinerea* WT, comparing three independent Bc-sRNA STTM T2 plant lines (#3, #4, #6) with *A. thaliana* WT and a non-sense STTM T2 plant line serving as negative controls. Lesion sizes were measured at 48 hpi and a minimum 15 lesions per plant line were used for statistical analysis using ANOVA followed by a Tukey post-hoc test with *p*-value threshold *p* < 0.05. The scale bars represent 1 cm. C) Quantitative reverse transcriptase PCR measuring mRNA levels of Bc-sRNA target genes in *A. thaliana AtMPK1, AtMPK2*, and *AtWAK* comparing STTM lines and WT during *B. cinerea* infection at 24 hpi. Lines in scatter plots represent the mean and the standard deviation. Each gene was measured in at least five biological replicates. *At-Actin* was used as a reference gene. *AtBAK1* and *AtPDF1*.*2* were measured as plant immunity marker genes induced during *B. cinerea* infection. Statistical analysis was performed using ANOVA followed by a Tukey post-hoc test with *p*-value threshold *p* < 0.05.

We previously found that expressing a triple STTM in *A. thaliana* that blocked small RNAs of the oomycete pathogen *Hyaloperonospora arabidopsidis* led to reduced disease levels (Dunker et al., 2020). We hypothesized that combining binding sites of different types of pathogen and parasite small RNAs that are known to induce cross-kingdom RNAi might create a multi-pathogen resistant plant genotype. We designed a “master” STTM to block the two *B. cinerea* Bc-sRNA3.2 and Bc-siRNA5, the two *H. arabidopsidis* small RNAs, Hpa-sRNA2 and Hpa-sRNA90 (Dunker *et al*., 2020), as well as the two *Cuscuta campestris* ccm-miR12497b and ccm-miR12480 (Shahid *et al*., 2018) and the *Phytophthora infestans* Pi-miRNA8788 (Hu *et al*., 2022). *A. thaliana* master STTM plants grew and developed normally and did not show any pleiotropic defect. We inoculated three independent T2 master STTM plant lines with *B. cinerea* and found that all transgenic lines exhibited reduced fungal lesion sizes (Figure S8A), confirming our results obtained with the Bc-sRNA STTM plants. Interestingly, a master STTM plant also revealed reduced disease levels when inoculated with the oomycete *H. arabidopsidis* (Figure S8C). Reduced infection was not due to constantly enhanced plant immunity, since inoculating Bc-sRNA- or master STTM plant lines with the bacterial pathogen *Pseudomonas syringae* DC3000 did not result in fewer reduced bacterial colonies (Figure S8D). Based on these observations, we propose that STTMs might be a future application against pathogen small RNAs to design multi-pathogen resistant plants.

## Discussion

In this study, we demonstrated that BcRDR1 is a pathogenicity factor that is required for cross-kingdom RNAi in the fungal plant pathogen *B. cinerea*. Previously, a *bcdcl1/2* mutant was characterized to be impaired in pathogenicity, Bc-sRNA biogenesis, and cross-kingdom RNAi (Wang *et al*., 2016, Weiberg *et al*., 2013). This *dcl1dcl2* ko mutant exhibited reduced growth in axenic culture as well as on plant leaves that could be partially complemented by ectopic Bc-sRNA expression in transgenic *A. thaliana* (Weiberg *et al*., 2013). Effective *BcDCL*1/*BcDCL2* gene knockdown via both host-induced gene silencing or RNA spray-induced gene silencing was sufficient to reduce *B. cinerea* disease symptoms on various plant species and tissues supporting a functional role of BcDCLs in pathogenicity (Qiao *et al*., 2021, Qiao *et al*., 2023, Wang *et al*., 2016). Recently, the impact of *bcdcl1dcl2* ko generated in the *ku70* ko background was debated to not exhibit any growth defect in axenic culture and no or only mildly reduced pathogenicity on plant leaves (He *et al*., 2023, Qiao *et al*., 2023).

The *bcrdr1* ko mutants characterized here did not display any growth defect when grown in axenic culture, but a significant reduction in lesion size formation during plant infection and reduced *in planta* fungal biomass when infecting the two plant host species *S. lycopersicum* and *A. thaliana*. Furthermore, *bcrdr1* ko mutants were impaired in activating a Bc-sRNAs-responsive switch-on cross-kingdom RNAi reporter expressed in transgenic *A. thaliana*. Moreover, we confirmed with further evidence the importance of cross-kingdom RNAi in *B. cinerea* pathogenicity using a STTM approach. Blocking Bc-sRNAs via STTM expression in *A. thaliana* led to reduced *B. cinerea* disease symptoms. A similar approach demonstrated the relevance of cross-kingdom RNAi in the oomycete pathogen *H. arabidopsidis* infecting *A. thaliana* (Dunker *et al*., 2020).

*RDR* knock out in different fungal plant pathogens resulted in pleiotropic defects on growth, development, and pathogenicity. The rice blast fungus *M. oryzae* comprises three RDRs named MoRDRP1-MoRDRP3, and the MoRDRP1 is the closest orthologue of BcRDR1. Isolated *mordrp1* ko mutants exhibited reduced growth, conidia formation, and disease symptoms on rice leaves (Raman *et al*., 2017). Unlike *bcrdr1, mordrp1* revealed only moderate changes in small RNA production, but *mordrp2* ko was strongly impaired in retrotransposon- and intergenic-derived 21-23 nt small RNA production. Thus, the underlying MoRDRP1 mode of action in pathogenicity remained obscure. In the head blight-inducing fungal pathogen *F. graminearum* five RDRs, named FgRdRP1-FgRdRP5, were identified. In two independent studies, distinct roles of FgRdRPs in fungal development and pathogenicity were reported (Chen *et al*., 2015, Gaffar *et al*., 2019). Studying *fgrdrp* ko mutants revealed neither altered scab disease symptoms when infecting flowering wheat heads or rot symptoms on tomato fruits, nor defects in vegetative development under normal or abiotic stress growth conditions (Chen *et al*., 2015). On the contrary, different *fgrdrp* ko mutants displayed alterations in asexual and sexual development. Moreover, infecting wheat spikes with *fgrdrp2, fgrdrp3* and *fgrdrp4* ko mutants at 9 days post inoculation resulted in reduced head blight symptoms, which correlated with lower pathogen DNA content and levels of the mycotoxin Deoxynivalenol per seed dry weight (Gaffar *et al*., 2019). The role of FgRdRPs in small RNA biogenesis has not been studied so far. The alternating numbers of RDR homologs in different fungal species and their distinct roles in small RNA biogenesis and pathogenicity indicate a rapid RDR neo-functionalization of the RNAi components in fungi. The unknown roles of BcRDR2 and BcRDR3 in *B. cinerea* need to be investigated in future studies.

Movement of small RNAs between fungi and plants has been observed multiple times (Wang & Dean, 2020). Cross-kingdom RNAi has been discovered in other plant- or animal-associated fungal, oomycete, and bacterial microbes, for pathogens and symbionts (Cai *et al*., 2019, Weiberg & Jin, 2015). It will be interesting to investigate whether microbial RDRs play a broader role in diverse plant-microbe interactions and in cross-kingdom RNA communication.

## Materials and methods

### Fungal and plant materials and growth conditions

*Botrytis cinerea* (Pers.: Fr.) strain B05.10 was used for this study. Standard cultivation was carried out on HA medium (10 g/L malt extract, 4 g/L yeast extract, 4 g/L glucose, 15 g/L agar). *B. cinerea bcrdr1* ko and *bcdcl1dcl2* ko mutant strains were grown on HA medium supplemented with 70 μg/mL hygromycin B (Carl Roth GmbH). Fungal plates were incubated at room temperature in a growth chamber under long wavelength UV light (EUROLITE; 20 W to stimulate sporulation. The tomato (*Solanum lycopersicum* (L.) cultivar Heinz) used in this study was grown under the condition of 24 °C, 16 h light/8 h dark, 60% humidity in a growth cabinet. *Arabidopsis thaliana* (L.) ecotype Col-0 was grown under short day condition (10 h light/ 14 h dark, 22°C, 60% relative humidity) for *Botrytis* infection. For *H. arabidopsidis* infection, *Arabidopsis* Col-0 was grown under long day condition (16 h light/ 8 h dark, 22°C, 60% relative humidity). Transgenic T1 and T2 *A. thaliana* lines were selected on 1/2 Murashige and Skoog (MS) medium supplemented with kanamycin (Carl Roth; 100 μg/mL). Independent plant transformant lines were indicated (e.g., STTM #3).

### Fungal and plant transformation

The Golden Gate cloning strategy was used for cloning fungal gene ko cassettes following the instruction, as described in Binder *et al*. (Binder *et al*., 2014). *B. cinerea* was transformed as previously described (Muller *et al*., 2018) with minor modifications. Transformed fungal protoplasts were mixed with SH agar (0.6 M sucrose, 5 mM Tris-HCl (pH 6.5), 1 mM (NH_4_)H_2_PO_4_, 8 g/L agar) without any antibiotics and incubated in darkness for 24 hours. A second layer of fresh SH agar containing hygromycin B (25 µg/ml) was added to the top after pre-incubation. The plates were further incubated in darkness until isolation of fungal transformants. For *A. thaliana* transformation, a previously described floral dip method was used (Clough & Bent, 1998).

### STTM and cross-kingdom RNAi reporter transgenic *A. thaliana*

The Golden Gate cloning strategy was used to clone plant expression vectors (Binder *et al*., 2014). The STTM sequences were designed, as described previously (Tang *et al*., 2012), and flanking region with BsaI recognition sites were introduced. A previously designed transgenic *A. thaliana* line that expressed a STTM with a randomized sequence of *H. arabidopsidis* small RNA target site (Dunker *et al*., 2020) was used as a non-sense STTM control in this study.

For *A. thaliana* reporter lines, a Csy4 coding sequence was synthesized by MWG Eurofins with codon optimization for expression in plants, and the reporter cassette were assembled, as previously described (Dunker *et al*., 2020). The native promoter of the Bc-sRNA3.1 target gene *AtPRXIIF* was used to control the transcription level of the *Csy4* transgene in order to mimic natural Bc-sRNA target expression.

### Plant infection assay

*S. lycopersicum* pathogenicity assays were performed on detached leaves from 4-to 5-week-old plants. *B. cinerea* conidia were resuspended in 1% malt extract at a final concentration of 5×10^4^ conidia/ml. *S. lycopersicum* leaves were inoculated with 20 μl conidia suspension and inoculated in a humidity plastic box. The detached leaves were placed on moist filter paper and then incubated in a closed plastic box. *A. thaliana* pathogenicity assays were performed on detached leaves from 5-week-old plants by inoculation with 2 × 10^5^ conidia/ml *B. cinerea* conidia resuspended in 1% malt extract. 15 μl conidia suspension was dropped at the center of each leaf and inoculated in a humidity plastic box. Infected leaves were photographed, and lesion area was measured using the Fiji software (ImageJ version 2.1.0/1.53c). For recording GFP cross-kingdom RNAi reporter time course, 3-week-old *A. thaliana* were cultivated on ½ MS + 1% sucrose agar plates and inoculated with 2×10^5^ conidia/ml *B. cinerea* resuspended in 1% malt extract medium pre-incubated for 1 hours. The pre-incubated conidia suspension was washed twice with sterile water before 5 µl was dropped at the center of one leaf per seedling. *Hyaloperonospora arabidopsidis* (GÄUM.) isolate Noco2 was maintained on *A. thaliana* Col-0 WT plants. Two-week-old plants were inoculated with 2×10^4^ conidia/ml suspension. Samples were harvested at 7 dpi into 10 ml of sterile water. The sporangiophore numbers were counted on detached cotyledons using a binocular.

*Pseudomonas syringae* pv. *tomato* DC3000 was streaked out on LB agar plates with Rifampicin for 2 days at 28 °C. A single colony from the plate was inoculated with LB liquid medium with Rifampicin overnight. *Pseudomonas* cells were harvested and re-suspended in 10 mM MgCl_2_ with 0.04% Silwet L-77 and adjusted to OD_600_ = 0.02. 5-week-old *A. thaliana* plants were sprayed with *Pseudomonas* suspension. Samples were harvested at 3 dpi and 4 leaf discs per plant were collected then homogenized in 10 mM MgCl_2_ for one biological replicate. A serial dilution on LB agar plates with Rifampicin was performed to count colony forming units.

### Genotyping PCR

A CTAB method followed by chloroform extraction and isopropanol precipitation was used for DNA extraction from fungal mycelium (Chen & Ronald, 1999). The GoTaq G2 Polymerase (Promega) was used for genotyping PCR using PCR primers as listed in Table S1.

### Quantitative PCR

A CTAB method was used for total RNA extraction (Bemm *et al*., 2016). Genomic DNA was removed by DNase I (Sigma-Aldrich) treatment following the manufacturer’s instruction. 1 μg of total RNA from each sample was used for cDNA synthesis using the SuperScriptIII reverse transcriptase (ThermoFisher Scientific). Gene expression was measured by quantitative real-time PCR using the Primaquant low ROX qPCR master mix (Steinbrenner, Laborsysteme). Differential expression level was calculated using the 2^-^ΔΔ^Ct^ method (Livak & Schmittgen, 2001). Primers used in quantitative PCR are listed in Table S1.

### Stem-loop reverse transcription PCR

Small RNA detection by stem-loop reverse transcriptase PCR was carried out following the protocol, as described (Varkonyi-Gasic *et al*., 2007). 1 μg of total RNA was used for small RNA-specific RT and PCR. Reverse transcriptase PCR products were separated on a 10% non-denaturing polyacrylamide gel followed by ethidium bromide staining. Primers used in stem-loop reverse transcriptase PCR are listed in Table S1.

### Microscopy

Single images of GFP signals were recorded on a Leica DM6 B upright microscope equipped with a Leica DFC9000 GT camera. For time course video, the Leica DMi8 Thunder Imager equipped with a Leica DFC9000 GT camera was used. Imaging was set to start 6 h post conidia inoculation and images were taken in cycles of 22.5 mins for 45 h in total. Raw imaging data were processed using the Leica LAS X microscope software. Video editing was performed using Adobe Premiere Pro CC version13.0.

### Small RNA sequencing

Small RNAs were isolated from total RNA extracts using 15% polyacrylamide gel electrophoresis. Isolated small RNAs were subjected for library cloning using the Next Small RNA Prep kit (NEB) and sequenced on an Illumina NextSeq 2000 platform. The Illumina sequencing data were analyzed using the GALAXY Biostar server (Giardine *et al*., 2005). Raw data were de-multiplexed (Illumina Demultiplex, Galaxy Version 1.0.0) and adapter sequences were removed (Clip adaptor sequence, Galaxy Version 1.0.0). Sequence raw data are deposited at the NCBI SRA server (BioProject ID PRJNA978613). Reads were then mapped to the reference genome assembly of *B. cinerea* (ASM14353v4) using the BOWTIE2 algorithm (Galaxy Version 2.3.4.2) in end-to-end alignment mode, without setting –k or –a options. To assign Bc-sRNA to different types of RNA encoding genes, the sequence information of *B. cinerea* ribosomal RNAs, transfer RNAs, small nuclear and nucleolar RNAs, and mRNA were downloaded from the Ensembl database. Retrotransposon sequences was used as annotated in a previous study (Porquier et al., 2021). Reads were counted and normalized on total *B. cinerea* reads per million (RPM).

### Phylogeny

CLC Main Workbench (version 20.0.4) was used for phylogenetic analysis and composing the DNA sequence labels. Alignment was conducted with amino acid sequences of each RDR by default multiple alignment algorithms. Phylogenetic tree was carried out by neighbor joining algorithm with 2000 replicates bootstrap.

### Data plotting and statistical analysis

GraphPad Prism 9 software was used for plotting and statistical analysis. For multi-samples comparison, one-way ANOVA with Tukey multiple comparisons test (p-value < 0.05) was carried out for data analysis. For *H. arabidopsidis* infection assay, unpaired t-test was carried out. Statistical significance was set for two-tailed p-value < 0.05 (*), p-value < 0.01 (**).

## Data accessibility

Sequencing data have been deposited in NCBI SRA (BioProject ID PRJNA978613).

## Acknowledgments

We thank Dr. Claude Becker for critical proofreading of this work. We want to thank the Gene Center Munich for Illumina NextSeq sequencing service. We would like to thank Dr. Martin Parniske for scientific discussions and providing access to the Golden Gate cloning system, Dr. Silke Robatzek and Dr. Eliana Mor for the access to and technical assistance with the DMi8 Thunder Imager microscope, and Dr. Dagmar Hann for sharing with us the *Pst* DC3000 strain. We thank Verena Klingl for technical support to isolate *Botrytis bcrdr1* ko transformants and Ignacio Mohr for helping with the *H. arabidopsidis* inoculation. We thank Franz Oberkofler for video editing. This work was supported by the German Research Foundation (DFG; Grant-ID WE 5707/2–1) in the framework of the Research Unit FOR5116. The funders had no role in study design, data collection and analysis, decision to publish or in preparation of the manuscript.

## Author contribution

APC and AW designed experiments and wrote the manuscript. APC conducted experiments. BL designed and cloned the STTM and conducted *A. thaliana* transformation. LH and CT designed *bcrdr1* ko vectors and performed fungal transformation. FD and LO designed and cloned the GFP cross-kingdom RNAi reporter and conducted *A. thaliana* transformation. FP conducted *B. cinerea* infection and stem-loop reverse transcriptase PCR assays. AW raised funding to conduct this project.

## Figure legends

Figure S1: Fungal RDR phylogenetic analysis (A) and amino acid sequence alignment of the RDR active site (B). The bar in (A) represents length of branch.

Figure S2: Expression levels of the three BcRDRs (A) and gene ko strategy of *BcRDR1* (B). Lines in scatter plots in A) represent the mean and the standard deviation. Statistical analysis was performed using ANOVA followed by a Tukey post-hoc test with *p*-value threshold *p* < 0.05.

Figure S3: Infection series and lesion measurement of *B. cinerea* WT and *bcrdr1* ko mutants on infected *S. lycopersicum* (A) and *A. thaliana* (B). The scale bars represent 1 cm. Lesion size induced by *B. cinerea* infection was measured at 48 hpi. Statistical analysis was performed using ANOVA followed by a Tukey post-hoc test with *p*-value threshold *p* < 0.05.

Figure S4: Mapping results of Bc-sRNAs obtained from *B. cinerea* WT and *bcrdr1* ko mutants at a *BcGyspy1* and *BcGypsy3* gene locus on chromosome 14 (A) and respective *BcGypsy1* and *BcGypsy3* mRNA levels (B). Lines in scatter plots represent the mean and the standard deviation. Statistical analysis was performed using ANOVA followed by a Tukey post-hoc test with *p*-value threshold *p* < 0.05.

Figure S5: Biological replicates of mRNA expression measurements of known Bc-sRNA target genes in *S. lycopersicum* (A) and *A. thaliana* (B) during infection with *B. cinerea* WT and *bcrdr1* ko mutants. Lines in scatter plots represent the mean and the standard deviation. Statistical analysis was performed using ANOVA followed by a Tukey post-hoc test with *p*-value threshold *p* < 0.05.

Figure S6: Independent infection series and fluorescence microscopy imaging using a GFP switch-on cross-kingdom RNAi reporter plant during infection with *B. cinerea* WT, *bcrdr1* ko mutants, and a *bcdcl1/2* mutant. *A. thaliana* WT plants were infected with *B. cinerea* WT to assess auto-fluorescence at infection sites, and non-infected GFP reporter plants were assessed for reporter auto-activity. The scale bars represent 500 µm.

Figure S7: Biological replicates of fluorescence microscopy imaging, mRNA and protein expression measurement with the GFP switch-on cross-kingdom RNAi reporter plant using 10x conidia concentration for *bcrdr1* inoculation. The red arrow in B) indicates the infection front of the *B. cinerea* inoculation. The scale bars in A) and C) represent 500 µm. Numbers in indicate GFP and RuBisCo signal intensities estimated by the FIJI software. Lines in scatter plots in C) and E) represent the mean and the standard deviation. Statistical analysis was performed using ANOVA followed by a Tukey post-hoc test with *p*-value threshold *p* < 0.05.

Figure S8: Infection series with *B. cinerea* WT (A), *H. arabidopsidis* (C) and *P. syringae* DC3000 (D) in transgenic T2 *A. thaliana* lines expressing a Bc-sRNA or master STTM. The scale bars in A) represent 1 cm. Lesion size induced by *B. cinerea* infection was measured at 48 hpi. B) mRNA expression levels of Bc-sRNA target genes *AtMPK1* and *AtWAK* in *A. thaliana*. Lines in scatter plots represent the mean and the standard deviation. Statistical analysis was performed using ANOVA followed by a Tukey post-hoc test with *p*-value threshold *p* < 0.05. C) *H. arabidopsidis* sporangiophores were counted at 7 dpi. *HpaEF1α* mRNA levels of *H. arabidopsidis* was measured at 4 dpi estimating the oomycete pathogen biomass. D) Colony-forming units (cfu) of *P. syringae* DC3000 were counted at 3 dpi. Statistical analysis in A), D) was performed using ANOVA followed by a Tukey post-hoc test with *p*-value threshold *p* < 0.05. Statistical analysis in C) was carried out by unpaired t-test with two-tailed p-value < 0.05 (*), p-value < 0.01 (**).

Video 1: Time course of the GFP cross-kingdom RNAi reporter activity upon *A. thaliana* leaf inoculation from 6-51 hpi with *B. cinerea* WT (left site) or *bcrdr1* ko mutant #2 (right site).

Table S1: DNA oligonucleotides used in this study.

