## Supplementary figures and images for "A fungal RNA-dependent RNA polymerase is a novel player in plant infection and cross-kingdom RNA interference"

### Supplementary figures

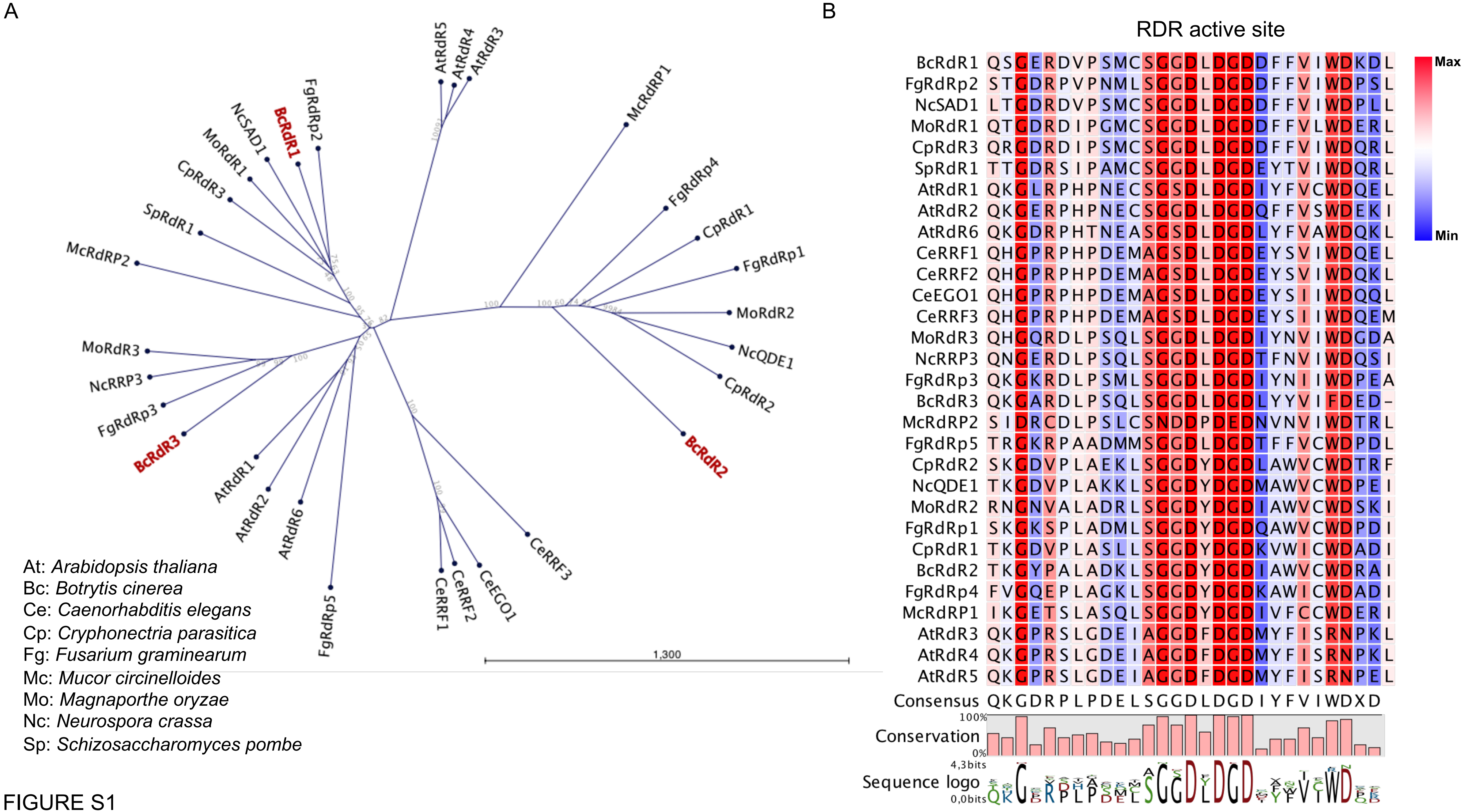

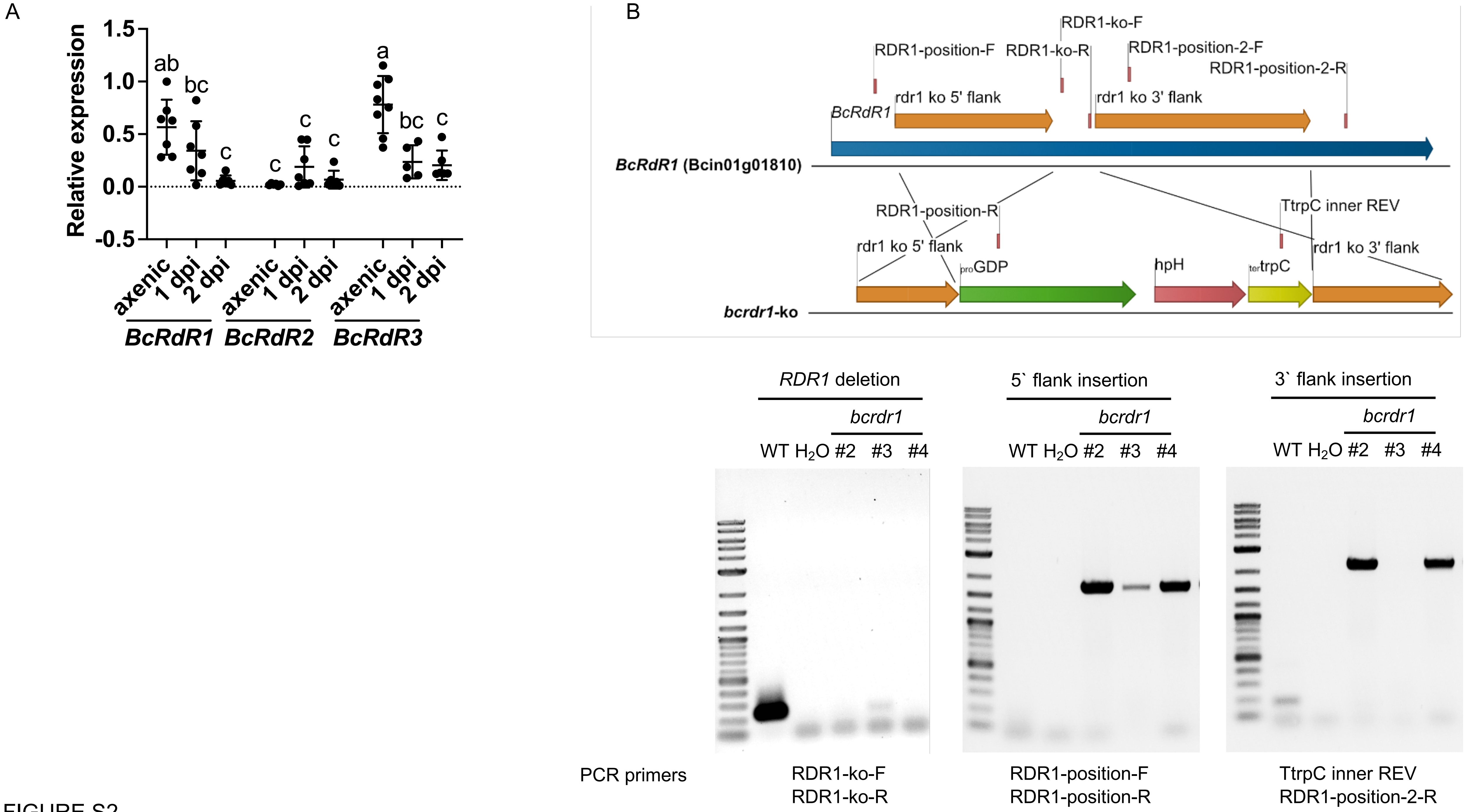

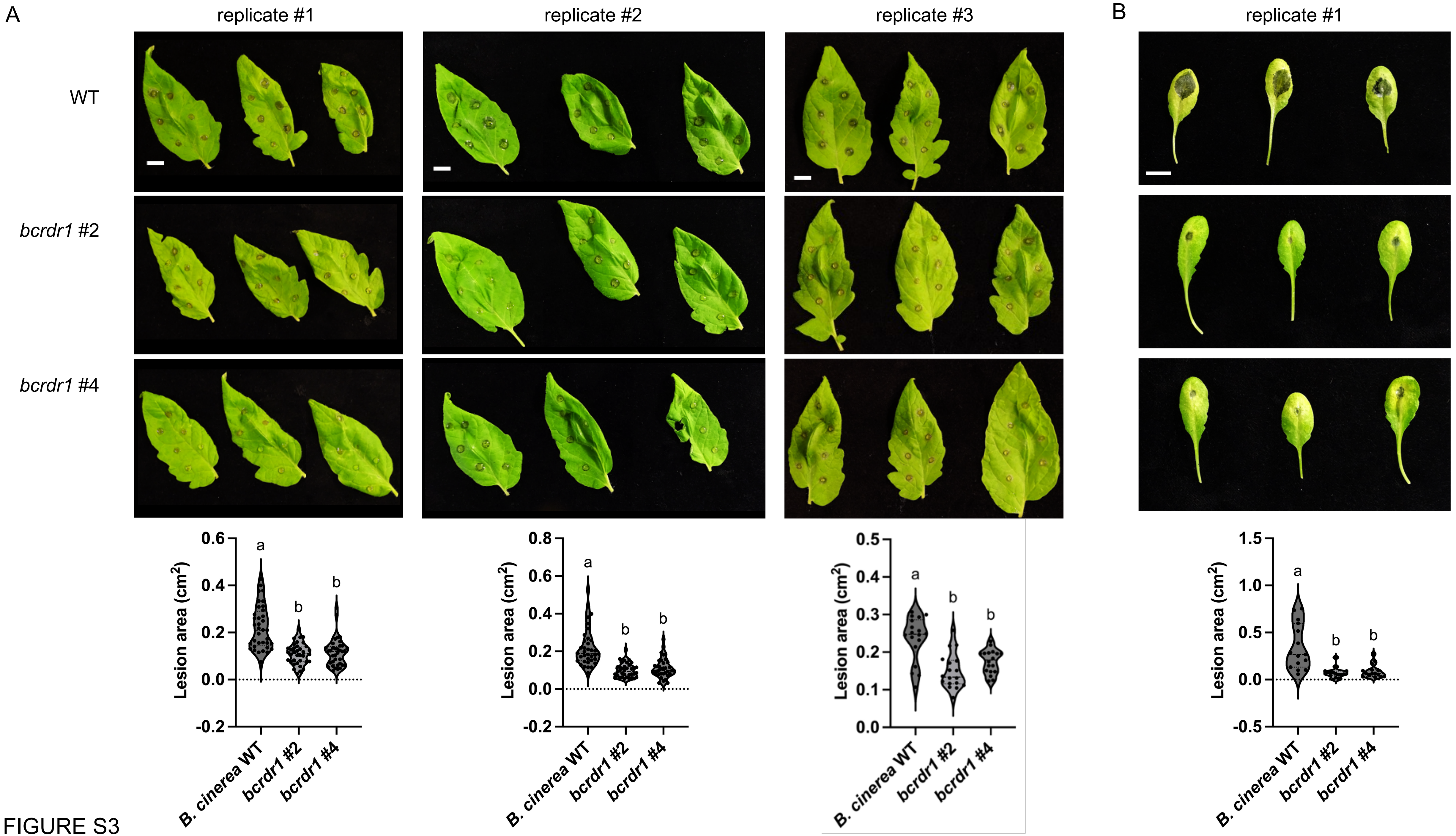

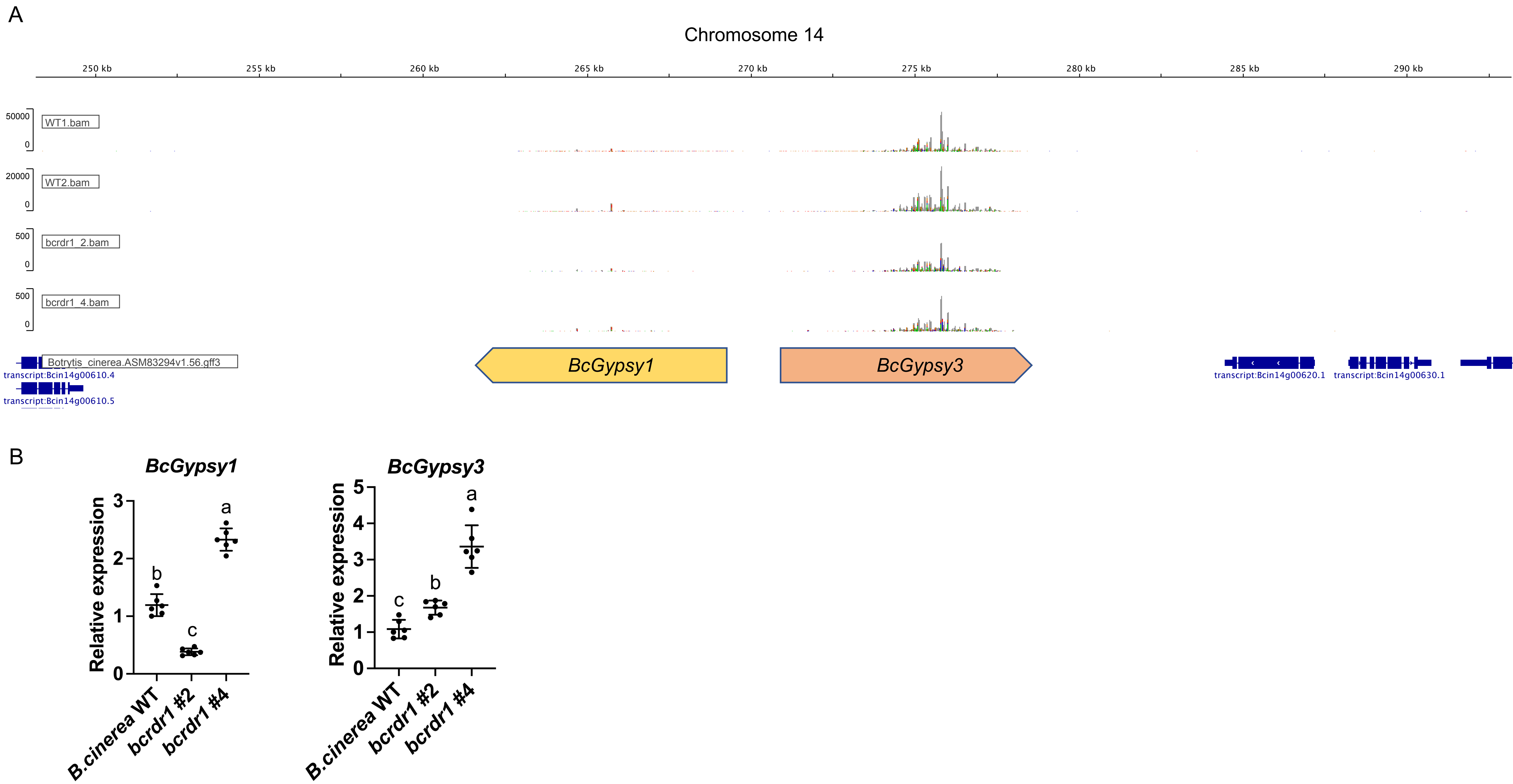

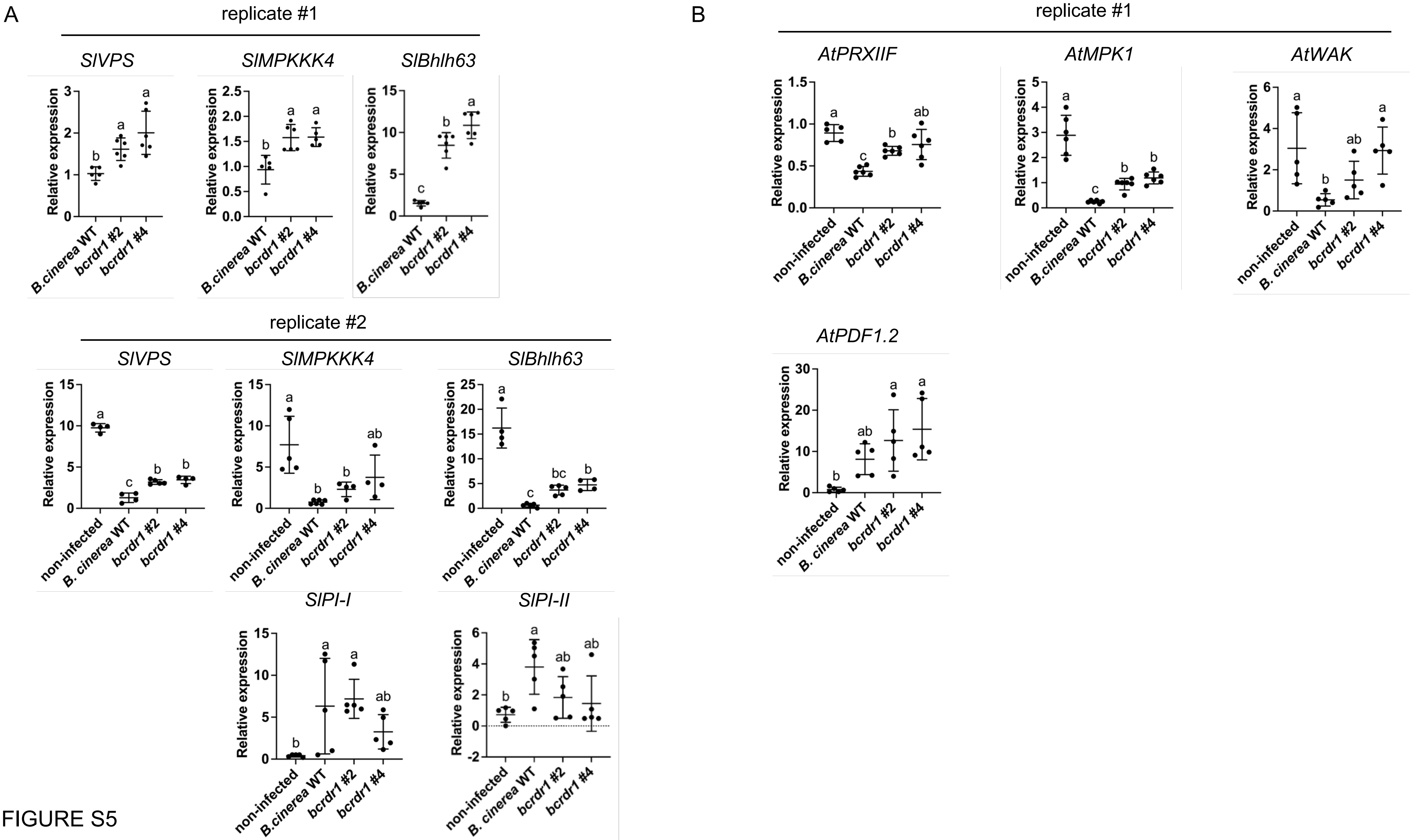

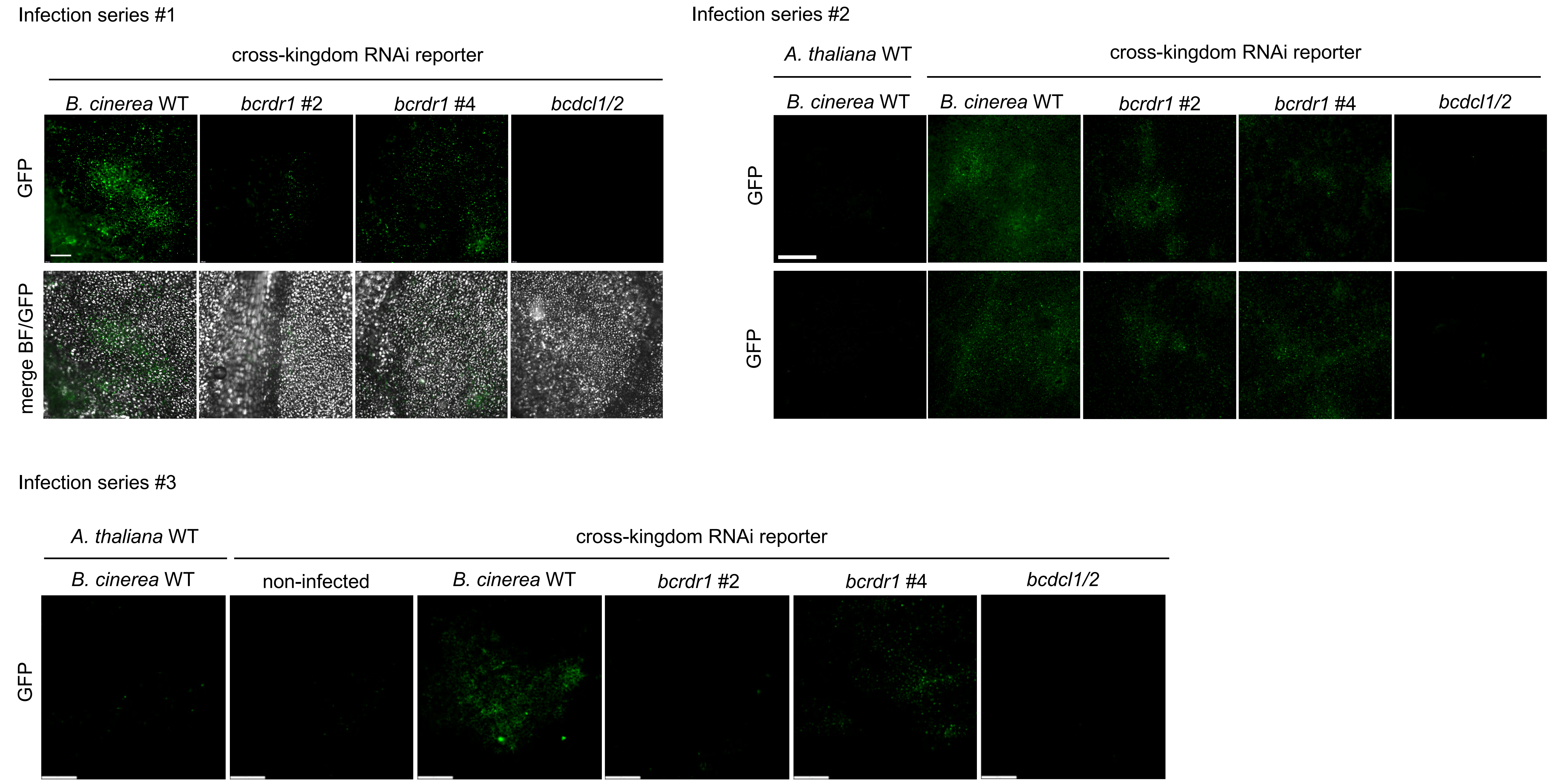

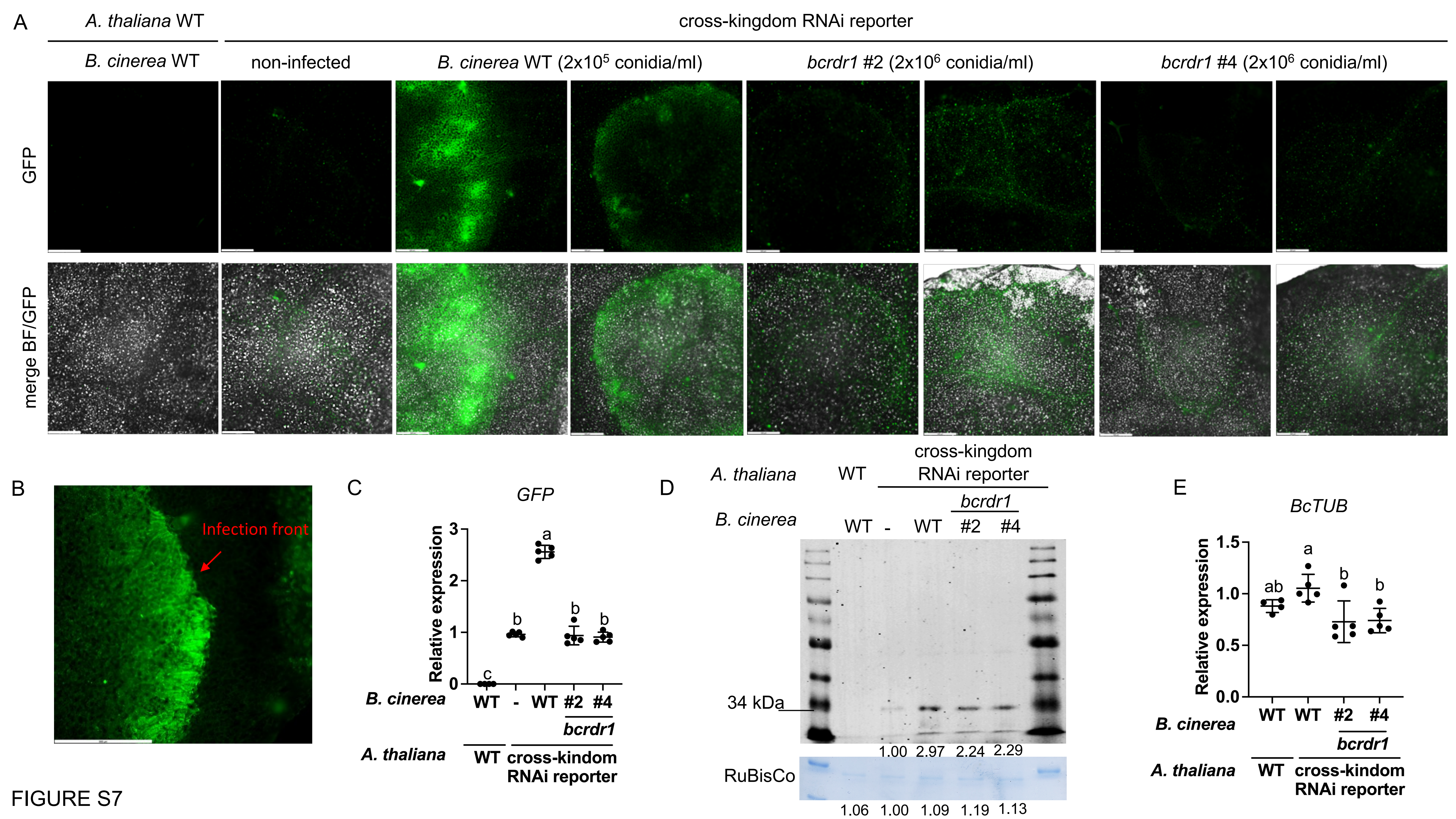

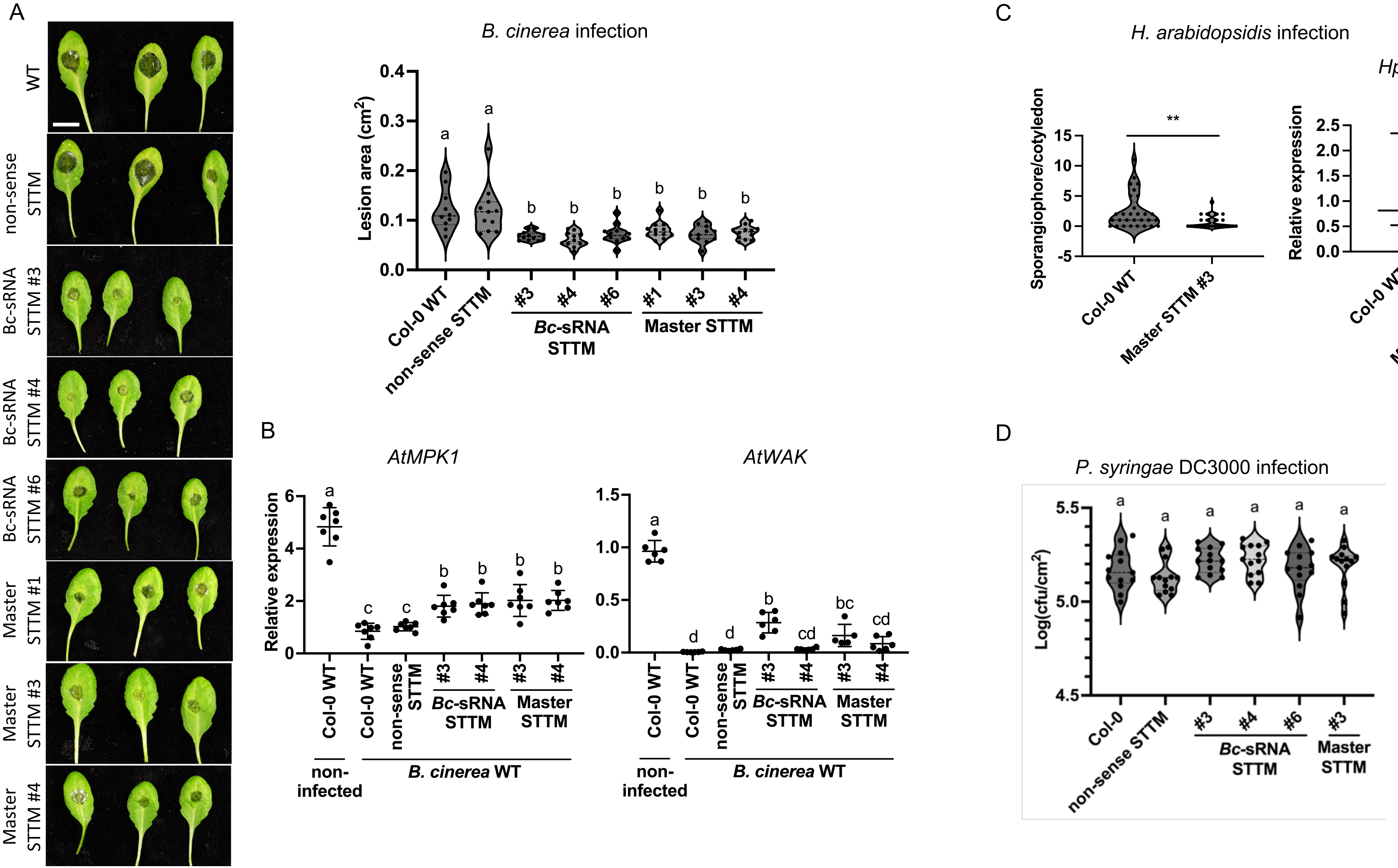
